# Global transcriptome analysis reveals *Salmonella* Typhimurium employs the nitrate-dependent anaerobic pathway to combat bile stress

**DOI:** 10.1101/2023.06.15.545048

**Authors:** Madhulika Singh, Deepti Chandra, Sirisha Jagdish, Dipankar Nandi

## Abstract

*Salmonella* Typhimurium is an enteric pathogen that is highly tolerant to bile. Next-generation mRNA sequencing was performed to analyse the stress and adaptive responses of *S*. Typhimurium to bile. We identified the cellular pathways affected during bile stress in wild type (WT) and a mutant lacking *csp*E (Δ*csp*E), which plays an essential role in protection from bile stress. We observed transcriptional upregulation of several genes involved in nitrate metabolism, in response to bile stress. These genes were also differentially expressed between the bile-resilient WT and the bile-sensitive Δ*csp*E strain. To understand the role of nitrate metabolism in bile stress response, we generated a strain lacking *fnr* (Δ*fnr*), which is the global regulator of nitrate metabolism in *S*. Typhimurium. *fnr* was highly induced in the bile treated WT strain but not in the Δ*csp*E strain. Notably, the Δ*fnr* strain was susceptible to bile-mediated killing. Our studies revealed a new role for *fnr* in mediating the bile stress response. In addition, a strain lacking *arc*A (Δ*arcA*), a two-component system response regulator involved in anaerobic metabolism, also showed a marked reduction in growth in presence of bile. This corroborated the significance of anaerobic metabolism in *S*. Typhimurium bile tolerance. Importantly, overexpression of *fnr* and *arc*A lowered reactive oxygen species and significantly enhanced the survival of the bile-sensitive Δ*csp*E strain. We also observed that *S*. Typhimurium pre-treated with nitrate displayed better growth in the presence of bile. Together, these results demonstrate that nitrate-dependent anaerobic metabolism promotes adaptation of *S*. Typhimurium to bile.

**Importance:** *Salmonella* Typhimurium, as an enteric pathogen, manifests an extreme example of bile tolerance. This study describes the diverse metabolic changes at the level of transcriptome in *S*. Typhimurium exposed to bile. We identified the differential expression of several genes involved in anaerobic metabolism between bile-tolerant WT and bile-sensitive Δ*csp*E strains. Two major regulators of anaerobic metabolism, *fnr* and *arc*A, support the growth of *S*. Typhimurium in bile. Our results highlight that, in presence of bile, *S*. Typhimurium activates genes involved in anaerobic metabolism, specifically nitrate metabolism, that improves survival of bacteria during bile stress.

## Introduction

Non-Typhoidal *Salmonella* (NTS) is a food and water-borne enteric pathogen associated with gastroenteritis in humans. The non-typhoidal serovars are responsible for 93 million enteric infections and 155,000 deaths due to diarrhea (1, 2). In immunocompromised individuals, the bacteria cause severe systemic infection leading to 680,000 fatalities each year (1, 3). In addition, the emergence of multi-drug resistant strains of *S. enterica* is a further challenge for global health (4). Overall, *S*. *enterica* has a significant contribution to morbidity and mortality world-wide (5, 6).

*S*. Typhimurium has a diverse set of stress responses that allow it to survive in hostile environments within the host. One of the important factors contributing to the successful colonization of *S*. Typhimurium in intestine is tolerance to high levels of bile. Bile acts as detergent to compromise bacterial cell membrane. In addition, it also exerts its antibacterial effect by compromising integrity of DNA, RNA and proteins, chelating metal ions such as calcium and iron (7–9). Bile also causes generation of reactive oxygen species, thereby causing oxidative damage to the cells (7, 10–12). As a pathogen, *S*. Typhimurum displays an extreme example of bile tolerance as it is able to colonize the hepatobiliary tract and gall bladder during infection (9, 13). Previously, our lab characterized Cold shock protein E (CspE) mediated bile stress response in *S*. Typhimurium (14). CspE is an RNA chaperone that binds to transcripts and increases their stability (15). *csp*E is indispensable for survival of *S*. Typhimurium in presence of bile as a *csp*E deletion strain (Δ*csp*E) is highly sensitive to bile stress (14). Besides, *csp*E and its homolog *csp*C also play a role in the pathogenicity of *S*. Typhimurium (15). Given the ability of *S*. Typhimurium to adapt to bile during infection, we decided to identify the pathways that differentiate the bile stress response of WT and *csp*E deletion strains, through whole genome transcriptome analysis. While RNA-Seq would capture a global, genome-wide transcriptional change in presence of bile stress, it cannot reveal a gene’s contribution to phenotype and survival in bile stress. Considering the essential role of *csp*E in mediating *S*. Typhimurium bile stress response, a comparison of transcriptome of bile treated WT and Δ*csp*E strains could lead to identification of genes important for survival in bile stress. Utilizing this approach, we identified the metabolic remodeling pathways that provides growth advantage to *S*. Typhimurium in the presence of bile.

*S*. Typhimurium possesses the ability to rapidly reprogram its metabolism in response to physiological stressors. Utilization of various carbon sources and respiratory electron acceptors in the inflamed intestine supports the growth of *S*. Typhimurium (16–18). The nitric oxide and superoxide generated by host immune response in gut to limit the growth of pathogen, can react in the intestine resulting in generation of nitrate (19, 20). However, *S*. Typhimurium can utilize nitrate as an alternative electron donor for anaerobic respiration (20, 21). This provides growth advantage to *S*. Typhimurium over the resident microbiota in gut that rely on fermentation, as nitrate is energetically superior to fermentation or other electron acceptors used during anaerobic respiration (20, 22, 23). Our findings provide insight into the importance of metabolic adaptation of *S*. Typhimurium to bile stress that requires nitrate and other anaerobic respiration. To the best of our knowledge, this is the first study to establish the roles of genes involved in nitrate and anaerobic metabolism, *fnr* and *arc*A, in combating bile stress in *S*. Typhimurium.

## Results

### RNA-Seq reveals major changes in gene expression of *S*. Typhimurium during bile stress

Global transcriptome profiles of WT and Δ*csp*E strains treated with 3% bile for 90 minutes were obtained (Fig. 1A). The principal component analysis suggested a higher variation between control and bile treated conditions than between the two strains (<u>Fig.S1A</u>). The transcriptional profiling of *S*. Typhimurium WT and Δ*csp*E strains revealed genes differentially expressed upon bile stress. A total of 466 genes were upregulated and 525 genes were downregulated in WT upon bile treatment. This constitutes 9.27% and 10.45% of the total 5,022 genes captured in the normalized expression data of RNA-Seq. As Δ*csp*E is sensitive to bile stress, a higher number of genes were differentially expressed compared to WT strain: 729 genes were upregulated whereas 834 genes showed downregulation in bile treated Δ*csp*E (Fig. 1B). Volcano plots depicted the same trend i.e., expression of higher number of genes was altered in Δ*csp*E compared to WT in presence of bile (<u>Fig. S1B</u>). Also, 305 upregulated and 302 downregulated genes were unique to Δ*csp*E upon bile treatment (Fig. 1C).

**Fig 1.**
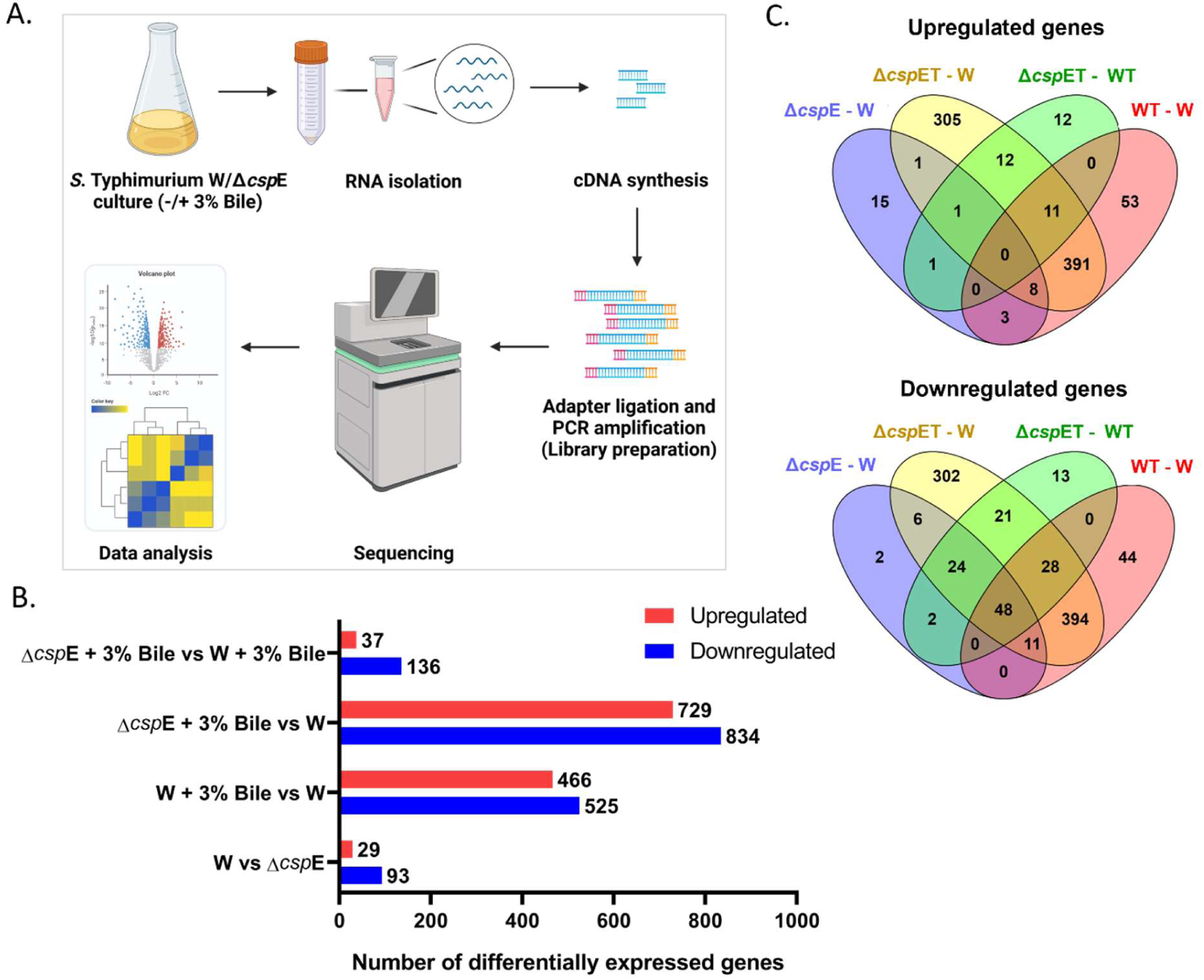
Visualization of changes in gene expression levels and comparison of upregulated and downregulated genes in response to bile treatment in *S*. Typhimurium WT and Δ*csp*E strains. A. Brief workflow of experiment. B. Histogram depicting distribution of upregulated and downregulated genes. C. Venn diagrams showing the number of genes that are >2 fold upregulated or downregulated. Conditions-Δ*csp*ET: Δ*csp*E + 3% Bile, WT: W + 3% Bile, Δ*csp*E: Δ*csp*E in LB, W: W in LB.

### The major alteration in transcriptome of bile treated *S*. Typhimurium involves changes in metabolism

The RNA-Seq data showed a distinct pattern of differentially expressed genes in WT and Δ*csp*E strains upon bile treatment (<u>Fig. S2</u>). To identify the major pathways that are affected in *S*. Typhimurium challenged with bile, a KEGG pathway analysis was performed with the differentially expressed genes in bile treated WT. The highest number of induced (144) as well as suppressed (59) transcripts were involved in metabolic pathways. Taken together, expression of 203 genes belonging to various metabolic pathways was significantly altered in presence of bile (Fig. 2A and B). Bile stress resulted in an overall induction of oxidative phosphorylation and citrate cycle (Fig. 2C). In addition, sulfur and nitrogen metabolism and glycolysis contributed to the anaerobic metabolism in the presence of bile (Fig. 2C). Further, we analyzed the differential expression profile of the transcripts to identify bile-specific and CspE-specific gene expression changes, particularly those involved in bacterial metabolism under bile stress. We observed that the transcripts of *glp* cluster, responsible for glycerol metabolism, were among the strongest induced in WT strain in presence of bile (Fig. 2D). While glpA, *glp*B and *glp*C constitute the operon that encodes anaerobic glycerol dehydrogenase, *glp*K encodes glycerol kinase (24, 25). However, the bile-sensitive Δ*csp*E strain showed an even better enrichment of these transcripts (Fig. 2D). Therefore, the induction of these genes in bile seems to be CspE-independent. A similar expression profile was observed for TCA cycle genes which showed CspE-independent upregulation in bile stress (<u>Fig. S3</u>).

**Fig 2.**
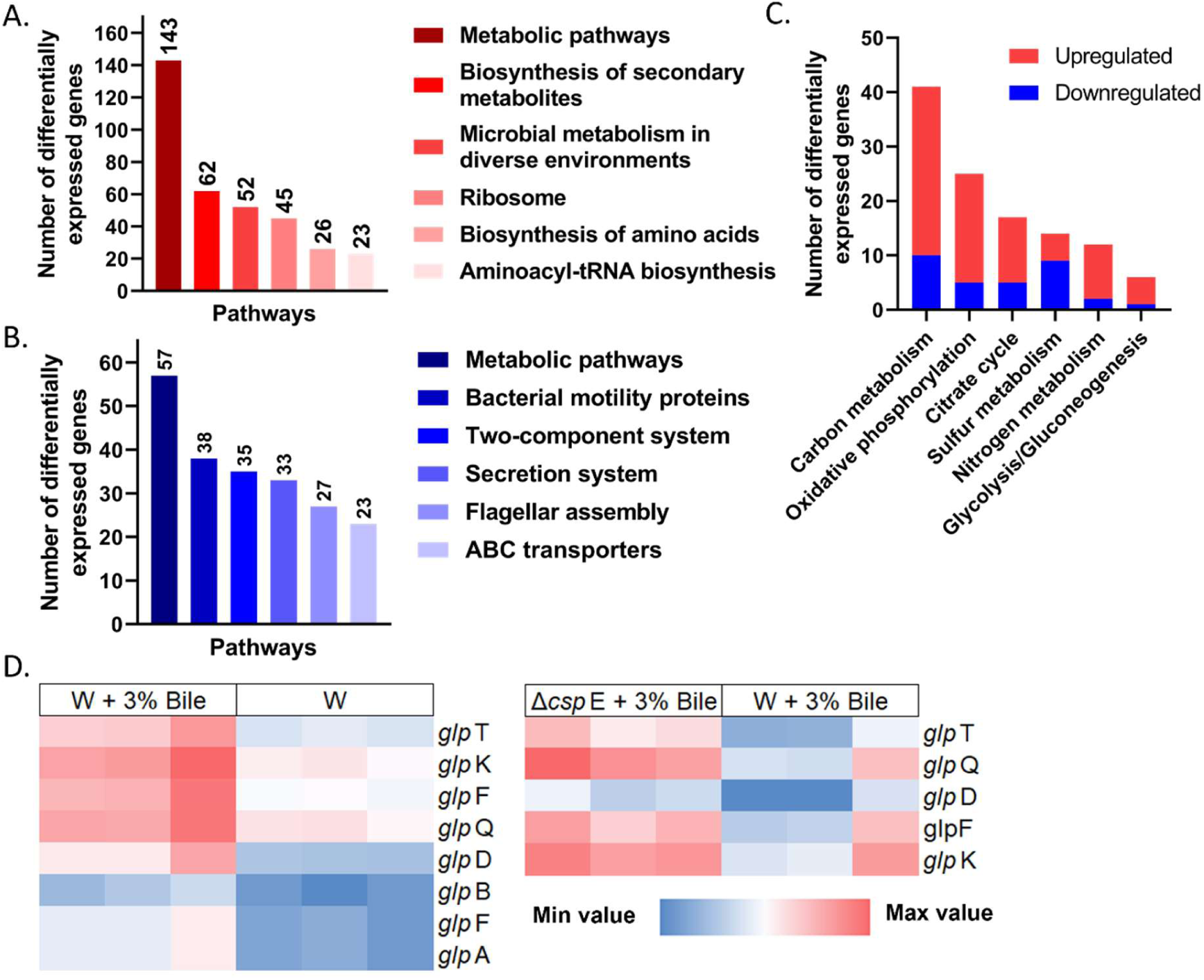
Transcriptome analysis of bile treated *S*. Typhimurium WT strain. Pathway analysis shows A. top most induced and B. suppressed pathways according to RNA-Seq. C. Histogram depicting modulation of various metabolic processes in response to bile. D. Heat map of *glp* genes with >1 Log_2_ fold increase represents induction of glycerol metabolism in bile-treated WT and Δ*csp*E strains.

Further, the twenty most enriched and depleted transcripts in bile treated WT and Δ*csp*E strains in RNA-seq are listed (<u>Table S6 and S7</u>). For validation, 5 of the 20 most upregulated and downregulated genes common to WT and Δ*csp*E were selected (<u>Fig.</u> <u>S4A and B</u>). Among the upregulated genes in RNA-seq, *nar*K is involved in nitrate metabolism while *cyo*A, *cyo*B and *cyo*D mediate aerobic metabolism. *ygb*L is involved in tartrate catabolic process (26). Corroborating the RNA-seq data, *nar*K, *ygb*L, *cyo*A, *cyo*B and *cyo*D were upregulated in bile treated WT and Δ*csp*E strains in q-PCR as well (<u>Fig. S4C</u>). *pdu*A, *yne*B, *omp*W, *yna*F and *yde*Y were significantly downregulated genes in RNA-seq. *pdu*A is a part of bacterial microcompartment for 1,2-propanediol degradation (27). Uniprot revealed that *yne*B is involved in degradation of autoinducer-2, a secreted signaling molecule for quorum sensing (28) whereas *yde*Y is a putative sugar transport protein for autoinducer-2. *omp*W is an outer membrane porin while *yna*F encodes a putative stress response protein. These genes are involved in bacterial stress responses. As observed in the RNA-seq, *pdu*A, *yne*B, *omp*W, *yna*F and *yde*Y showed downregulation in bile treated WT and Δ*csp*E in q-PCR, validating the RNA-seq data (<u>Fig. S4D</u>). Therefore, among the genes with the most altered expression, *nar*K, *cyo*A, *cyo*B, *cyo*D, *ygb*L and *pdu*A are involved in various metabolic processes whereas *yne*B and *yde*Y play a role in quorum sensing.

### Transcriptional analysis suggests nitrate metabolism is important for bile stress response and is compromised in bile sensitive Δ*csp*E

The RNA-seq analysis suggested that the nitrate and nitrite reductases are key differentially expressed gene clusters in *S*. Typhumurium bile stress response. The transcripts of nitrate metabolism genes were generally enriched upon bile treatment in in WT (Fig. 3A). NarL is the response regulator of the NarX-NarL two-component system that induces the transcription of *nar*GHJI, *nar*K in presence of high levels of nitrate (29–31). We validated the RNA-seq results by q-RT PCR and found that the response regulator *nar*L, the *nar*G, *nar*H and *nar*I encoding nitrate reductase A, were significantly upregulated in bile stress in *S*. Typhimurium WT strain. However, these genes showed a significant downregulation in Δ*csp*E strain compared to the WT strain upon bile treatment (Fig. 3B). In addition, nitrite reductase *nir*D was also induced upon bile treatment. *nrf*B, a pentahaem electron-transfer gene involved in reduction of nitrite/nitric oxide to ammonia was downregulated in bile treated WT. In Δ*csp*E its transcription was further reduced in presence of bile (Fig. 3C). Noticeably, the nitrate reductase response regulator transcribed in presence of low levels of nitrate, *nar*P was suppressed in bile treated WT even as the periplasmic nitrate reductase component *nap*H showed induction (<u>Fig. S5</u>). Besides nitrate metabolism, *S*. Typhimurium has several other types of anaerobic metabolism dependent on utilization of dimethyl sulfoxide, ethanolamine, thiosulfate (19, 32). While anaerobic metabolism supports the growth of the pathogen in inflamed intestinal lumen, it has been observed that activation of nitrate metabolism suppresses anaerobic metabolism dependent on other energetically inferior electron acceptors (22, 29, 33). We analyzed the RNA-seq data to understand the effect of bile stress on anaerobic metabolism dependent on other electron acceptors. Interestingly, the anaerobic respiration utilizing dimethyl sulfoxide (*dms*B, *dms*C) and ethanolamine (*eut*A, *eut*B, *eut*M, *eut*P) was downregulated upon bile treatment (Fig. 3D). 1.2-propanediol dependent fermentation was also suppressed in the presence of bile (*pdu*A, *pdu*F, *pdu*J, *pdu*K) (Fig. 3D). Notably, in presence of nitrate or nitrite, growth of Δ*csp*E was similar to WT (<u>Fig. S6 A and B</u>).

**Fig 3.**
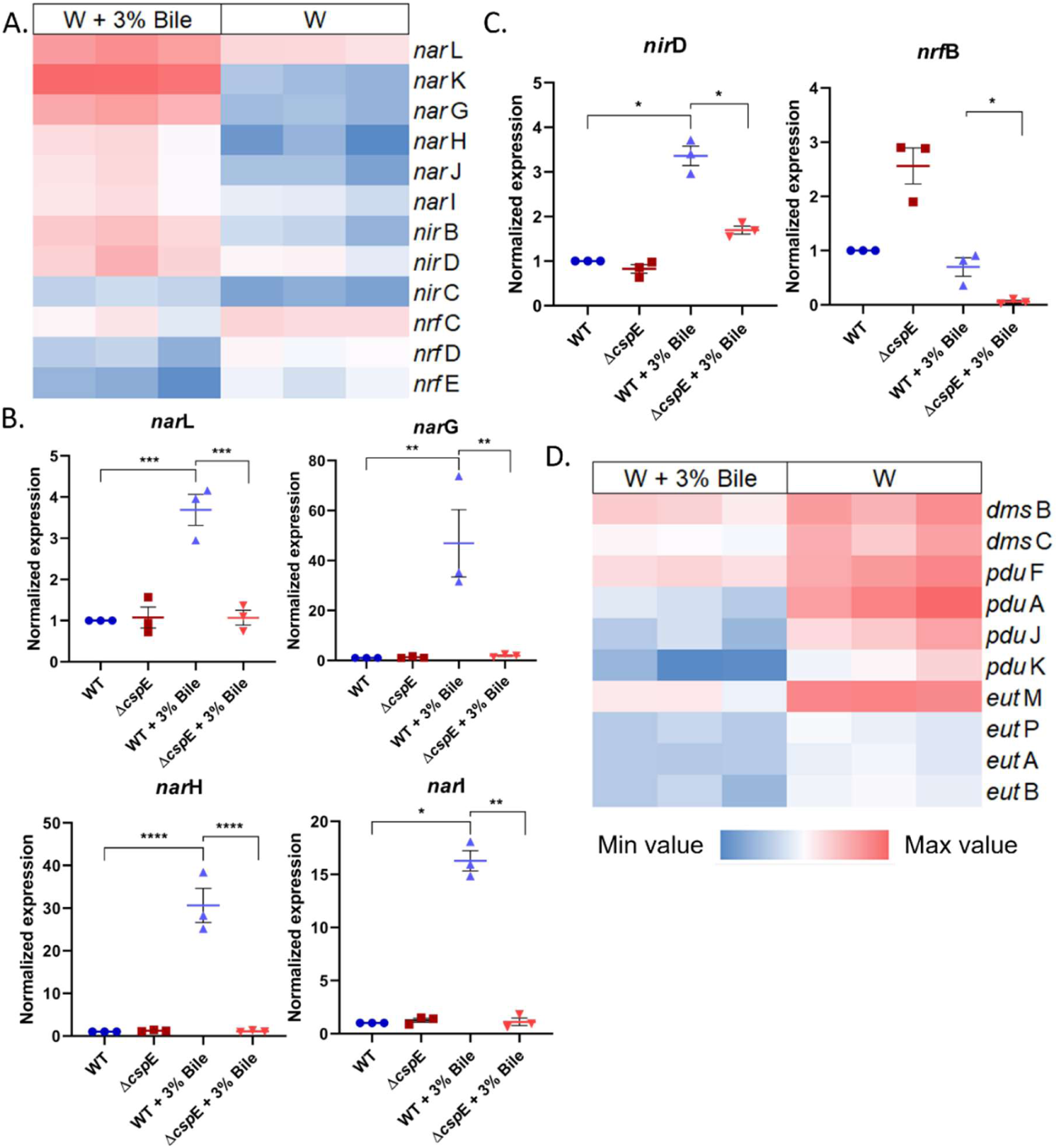
Transcriptional profile of genes involved in nitrate and anaerobic metabolism in bile treated *S*. Typhimurium. A. Heat map depicting differential expression of genes for nitrate metabolism between bile treated and untreated WT strain. B. qRT-PCR analysis of genes involved in nitrate utilization; nitrate response regulator gene *nar*L, nitrate reductases *nar*G, *nar*H and *nar*I. C. qRT-PCR of genes encoding nitrite reductase *nir*D and the periplasmic nitrite reductase *nrf*B. D. Heat map depicting differential expression of genes involved in anaerobic metabolism between bile treated and untreated WT strain. For qRT-PCR, data is shown as mean±SEM and is representative of 3 independent experiments. P values were measured by one-way ANOVA using Sidak’s multiple comparisons test. *p<0.05, **p<0.01, ***p<0.001, ****p<0.0001.

### *fnr* is required for growth of *S*. Tyhimurium in presence of bile

Nitrate reductases are involved in processes that generate energy for anaerobic metabolism utilizing nitrate as the electron acceptor. To determine the role of nitrate metabolism in bile stress response of *S*. Typhimurium, we generated a *nar*L deletion strain (Δ*nar*L). The Δ*nar*L strain showed similar growth to the WT strain in presence of bile (Fig. 4A). The similar phenotype of Δ*nar*L and WT strain in bile stress is possibly due to presence of other nitrate reductase response regulator such as *nar*P and nitrate reductase A isozyme NarZ (34). Fnr is the global regulator of nitrate/nitrite reductases in *S*. Typhimurium (33). Therefore, we hypothesized that targeting *fnr* would resolve the concern of functional redundancy associated with *nar*L, allowing us to study the role of nitrate metabolism in bile stress. To test this hypothesis, we first determined the expression pattern of *fnr* in bile stress. Upon bile treatment, *fnr* showed a slight but significant upregulation. Also, in presence of bile, it was downregulated in Δ*csp*E strain compared to WT (Fig. 4B). Subsequently, to confirm the biological function of *fnr* as a bile stress response gene, a *fnr* deletion strain (Δ*fnr*) was generated. The Δ*fnr* strain was sensitive to bile stress (Fig. 4C and D) and complementation of Δ*fnr* with p*fnr* enhanced its growth in presence of bile, similar to WT (Fig. 4E), demonstrating importance of *fnr* in mitigating bile stress.

**Fig 4.**
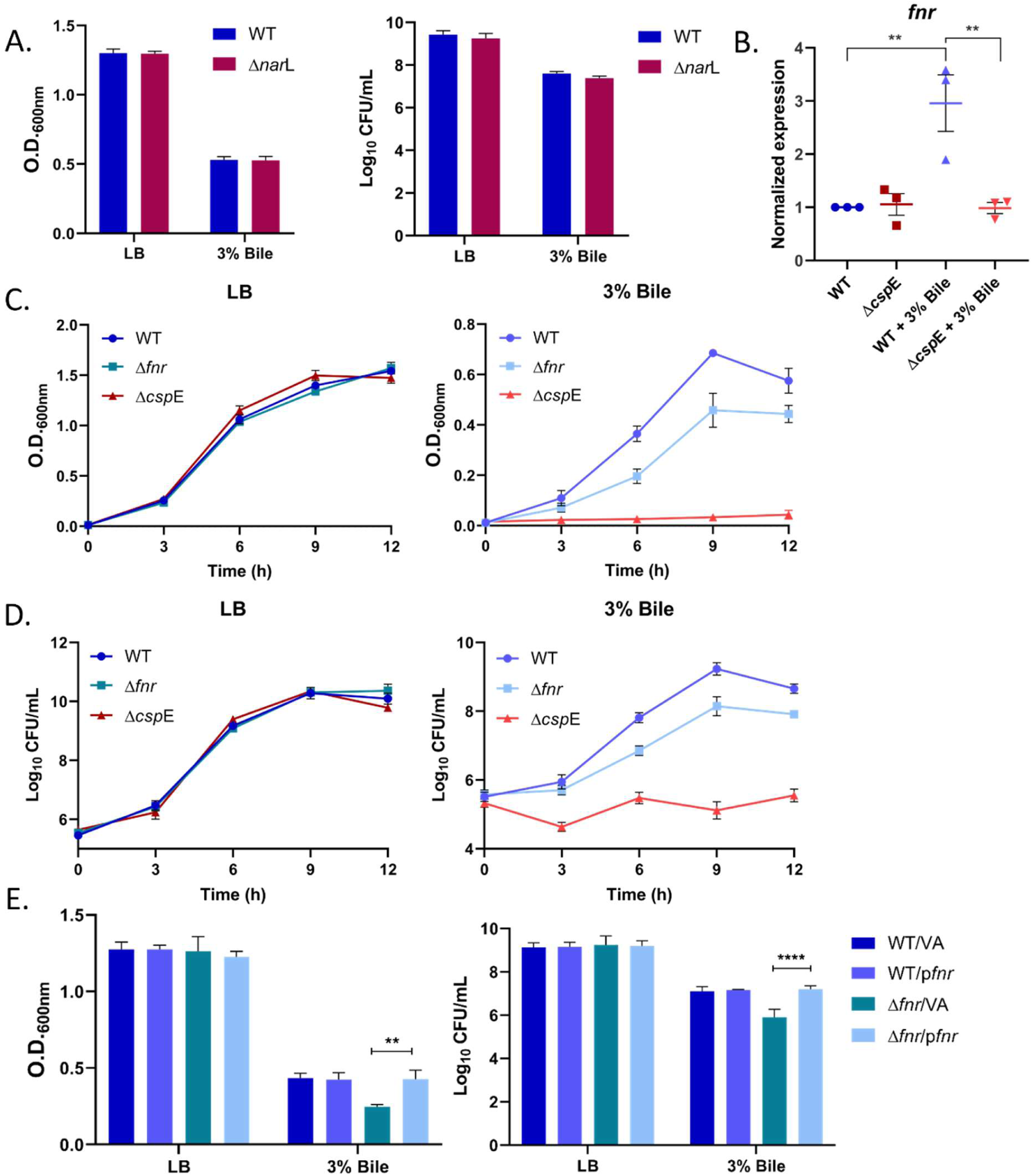
Role of *fnr* in *S.* Typhimurium challenged with bile. A. Growth of WT and Δ*nar*L strains in absence or presence of 3% bile. B. qRT-PCR analysis of *fnr* gene in bile treated WT and Δ*csp*E. C. Growth of WT, Δ*fnr* and Δ*csp*E strains with or without bile treatment. O.D. at 600 nm was measured at an interval of 3 h for 12 h. D. Cells were plated at appropriate dilutions and CFU were counted. E. Complementation of Δ*fnr* with p*fnr* in presence of 3% bile. Data shown as mean±SEM and is representative of 4 independent experiments. P values were measured by two-way ANOVA using Sidak’s multiple comparisons test. **p<0.01, ****p<0.0001.

### Anaerobic metabolism modulates the *S*. Typhimurium bile stress response

Although nitrate-independent anaerobic metabolism was suppressed upon bile treatment in WT, we found that several genes involved were further transcriptionally repressed in bile-sensitive Δ*csp*E compared to WT (<u>Fig. S7</u>). Therefore, we investigated the roles of the two-component signaling system response regulator *arc*A during bile stress response of *S*. Typhimurium (35). The q-PCR analysis revealed downregulation of *arc*A in bile-sensitive Δ*csp*E strain compared to WT (Fig. 5A). In order to target one of the key regulators of anaerobic metabolism in *S*. Typhimurium, we generated an *arc*A gene knockout strain (Δ*arc*A). The Δ*arc*A strain showed compromised growth in presence of bile supporting the role of anaerobic metabolism in bile tolerance of *S*. Typhimurium (Fig. 5B and C). Complementation of Δ*arc*A with p*arc*A restored the growth of Δ*arc*A strain similar to WT in presence of bile (Fig. 5D). These results establish a functional role of ArcA during bile stress.

**Fig 5.**
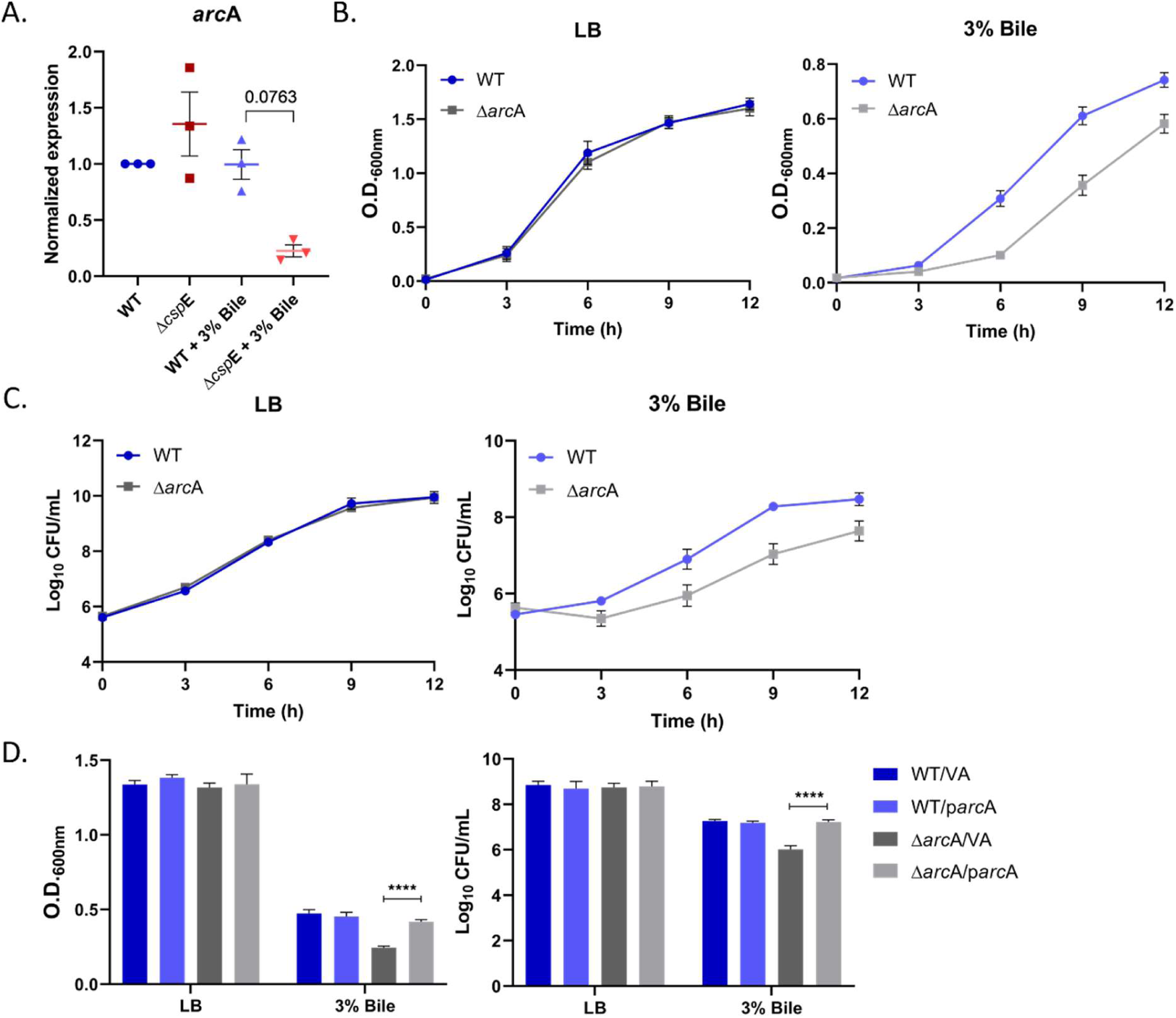
Functional significance of *arc*A in *S*. Typhimurium bile stress response. A. qRT-PCR analysis of arcA, 2 h post bile treatment. B. Growth of WT and ΔarcA strains with or without bile treatment was monitored kinetically at 3 h interval by measuring O.D. at 600 nm. C. Appropriate dilutions were plated and CFU was calculated. D. Complementation of ΔarcA with parcA in presence of bile. Data shown as mean±SEM and is representative of 4 independent experiments. P values were measured by two-way ANOVA using Sidak’s multiple comparisons test. ****p<0.0001.

### Overexpression of *fnr* and *arc*A enhance the survival of bile-sensitive Δ*csp*E strain by reducing intracellular ROS

The *S*. Typhimurium Δ*csp*E strain shows hypersensitivity to bile stress (14). *fnr* and *arc*A showed reduced expression in bile treated Δ*csp*E compared to WT suggesting that CspE is probably required for optimal expression of these genes. Therefore, we complemented the Δ*csp*E strain with each of these genes. There was no change in growth of WT strain overexpressing *fnr* and *arc*A upon bile treatment. However, the Δ*csp*E strain expressing *fnr* and *arc*A showed significant increase in survival in presence of bile. While expression of *fnr* rescued the growth of Δ*csp*E similar to that of WT (Fig. 6A), expression of *arc*A also led to partial but significant increase in growth in bile (Fig. 6B).

**Fig 6.**
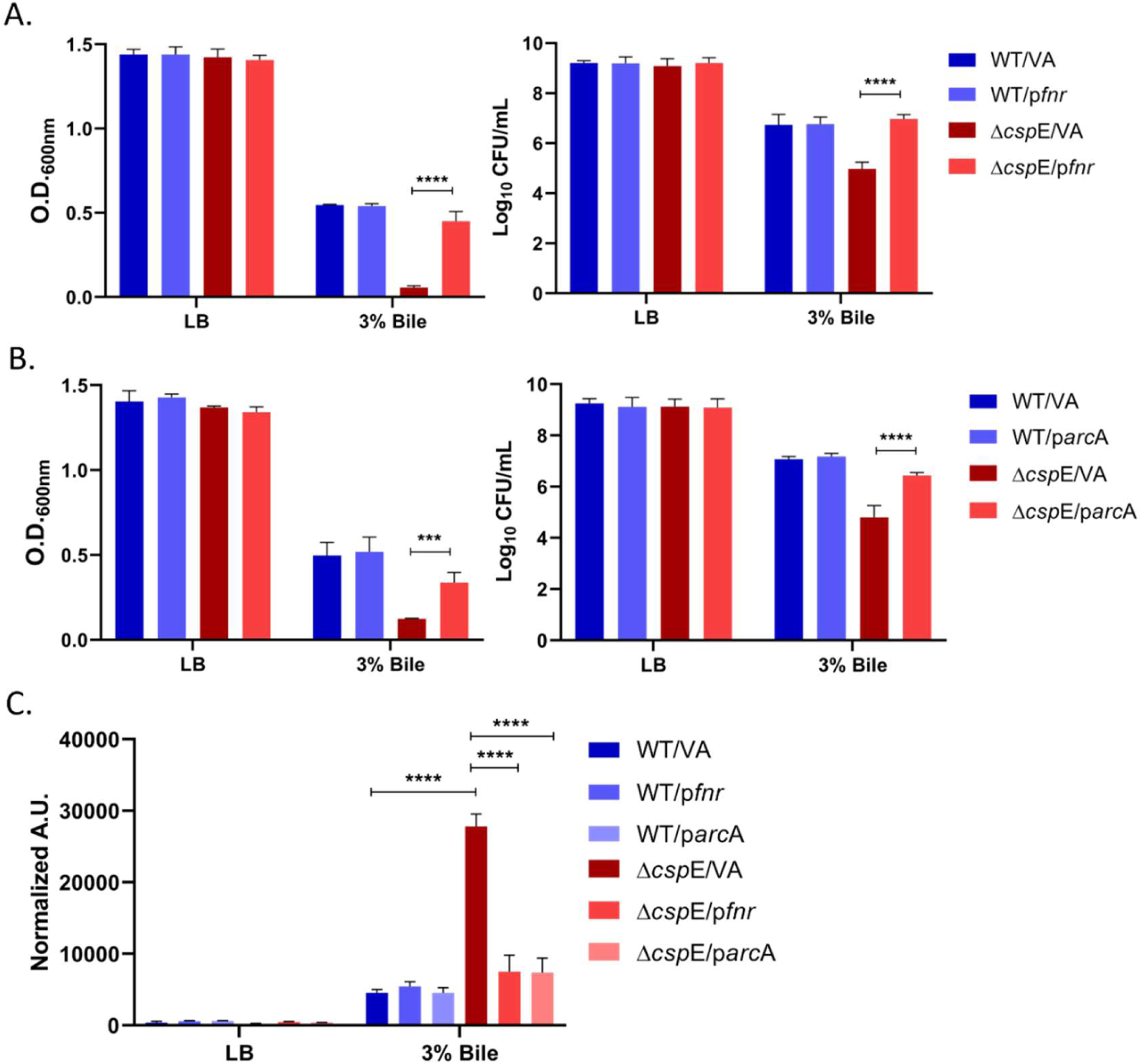
Growth of WT and Δ*csp*E strains expressing *fnr* and *arc*A in bile. Cells were treated with 3% bile for 6 h. O.D. at 600 nm was measured and CFU was calculated for indicated WT and Δ*csp*E strains. A. Overexpression of *fnr* in WT and Δ*csp*E strains. B. Overexpression of *arc*A in WT and Δ*csp*E strains. C. Estimation of intracellular ROS levels upon 3% bile treatment in WT and Δ*csp*E strains overexpressing *fnr* and *arc*A using DCFDA staining. Fluorescence intensity was measured using excitation and emission wavelengths of 485 nm and 535 nm respectively. Data shown as mean±SEM and is representative of 4 independent experiments. For ROS quantitation, data is shown as mean±SD and is representative of 3 independent experiments. P values were measured by two-way ANOVA using Sidak’s multiple comparisons test. ***p<0.001, ****p<0.0001.

To understand the mechanism by which *fnr* and *arc*A rescued bile-sensitive Δ*csp*E strain, we quantitated the level of intracellular reactive oxygen species (ROS) as *fnr* and *arc*A have been reported to protect the cells against oxidative stress (35, 36). Consistent with the previous studies (7, 10–12), bile treatment led to an increase in intracellular ROS in the WT strain (Fig. 6C). The increase in ROS was substantially higher in the Δ*csp*E strain which significantly reduced upon overexpression of *fnr* and *arc*A (Fig. 6C). These results suggest that expression of genes involved in nitrate and anaerobic metabolism alleviates the severe growth attenuation of Δ*csp*E strain in bile by reducing ROS.

### Pre-treatment with nitrate boosts the growth of *S*. Typhimurium WT strain in the presence of bile

The transcriptomic data suggested substantial induction of genes involved in nitrate metabolism in *S*. Typhimurium in response to bile. To understand whether nitrate metabolism provides a survival advantage in presence of bile, we pre-treated *S*. Typhimurium with nitrate before challenging with bile. Pre-treatment with sodium nitrate (NaNO_3_) significantly enhanced the growth of WT strain in bile stress (Fig 7A and B). The Δ*csp*E strain pre-treated with NaNO_3_ also showed slight increase in growth when exposed to bile (Fig 7A and B). Our results therefore demonstrate that *S*. Typhimurium grown in presence of nitrate shows better adaptation to bile stress.

**Fig 7.**
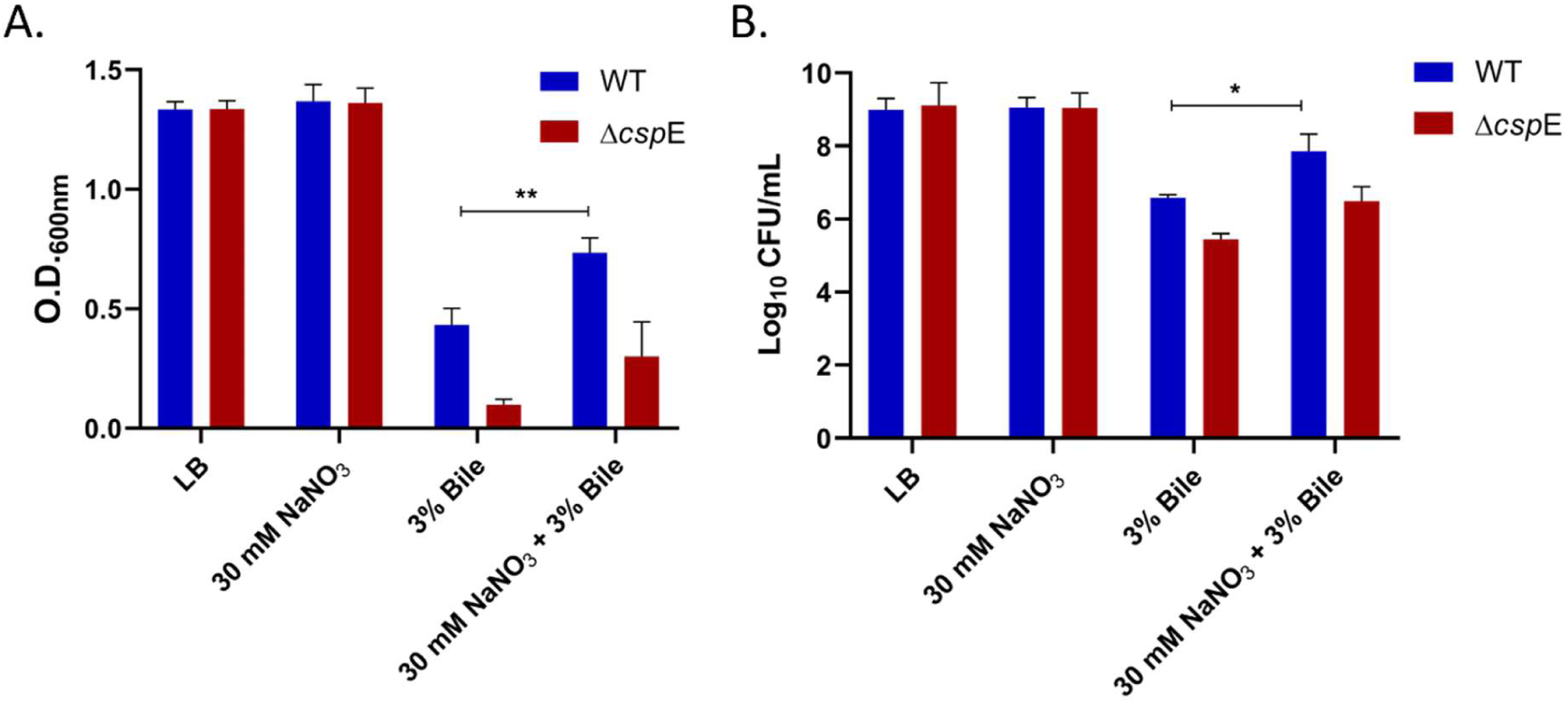
Effect of nitrate pre-treatment on growth of WT and Δ*csp*E strains in bile. A. O.D. and B. CFU of nitrate pre-treated cells after 6 hours of exposure to 3% bile. Data shown as mean±SEM and is representative of 4 independent experiments. P values were measured by two-way ANOVA using Sidak’s multiple comparisons test. *p<0.05, **p<0.01.

## Discussion

The bile resistance in *S*. Typhimurium is mediated by a myriad of stress response genes and is crucial for infection and survival within the host (9). Previously, our laboratory has shown that cold shock protein E (CspE) is essential for survival of *S*. Typhimurium under bile stress (14). In this study, RNA sequencing was performed to obtain the global transcriptome profile of *S*. Typhimurium WT and Δ*csp*E strains post bile treatment. While the WT strain can tolerate and grow in presence of bile, the Δ*csp*E strain is hypersensitive and shows severe growth attenuation in presence of bile. Initial analysis of differential expression profile and biological pathways revealed that *S*. Typhimurium challenged with bile undergoes major metabolic remodeling.

In addition to bile, *S*. Typhimurium also adapts to fluctuations in levels of oxygen during infection within the gastrointestinal tract of host. *S*. Typhimurium responds to the limited levels of oxygen in the gastrointestinal tract by utilizing different electron acceptors such as nitrate, dimethyl sulfoxide, tetrathionate etc. Our RNA-Seq data revealed that in *S*. Typhimurium, bile stress results in a metabolic switch, leading to induction of genes involved in anaerobiosis (Fig. 8). This includes enrichment of several transcripts responsible for nitrate metabolism, particularly nitrate reductase A encoded by *nar*GHJI. While *nar*GHJI is activated by nitrate, the cytoplasmic nitrite reductase *nir*BDC is activated by both nitrate and nitrite. On the contrary, the periplasmic nitrite reductase encoded by *nrf*ABCDEFG is repressed by nitrate and activated by nitrite (38). The induction of nitrate reductase A and nitrite reductase genes whereas inhibition of periplasmic nitrite reductase genes upon bile treatment indicates that the metabolic response is mainly nitrate-dependent. Unlike some obligate anaerobic bacteria that lack nitrate reductase activity, *S*. Typhiumurium expresses three nitrate reductases encoded by the *nar*KGHJI, *nar*UZYWV and *nap*FDAGHBC operons (19) which boosts the growth of *S*. Typhimurium in the intestinal lumen (22). Interestingly, the bile treated Δ*csp*E strain showed a compromised nitrate metabolism compared to WT at the transcript level. However, deletion of *nar*L, the two-component response regulator for nitrate reductase A, had no effect on survival of *S*. Typhimurium in presence of bile (Fig. 4). This may be attributed to the presence of other genes such as *nar*P and *nar*Z which would make the utilization of nitrate possible, even in absence of *nar*L. Therefore, we constructed a gene deletion mutant of *fnr* (fumarate nitrate reduction) which is a global transcription factor that regulates alterations in expression of genes that are important for survival of the enteric bacteria in low oxygen conditions (33). Fnr regulates expression of several anaerobic enzymes such as nitrate reductase A, cytoplasmic nitrite reductase, periplasmic nitrite reductase and fumarate reductase (38). Therefore, deletion of *fnr* would most likely cause major perturbance in anaerobic metabolism. Noticeably, bile stress resulted in induction of *fnr*. We also found that the isogenic strain of *S*. Typhimurium lacking *fnr* is susceptible to bile mediated killing (Fig. 4). Fnr has been associated with negative regulation of *pdu* operon as it directly binds to the promoter region of the operon, thereby suppressing propanediol utilization (39). We observed nitrate/nitrite metabolism to be induced whereas the propanediol operon was suppressed in presence of bile, implying an *fnr*-mediated response. However, the anaerobic dimethyl sulfoxide reductases (*dms*A, *dms*B and *dms*C), positively regulated by *fnr*, were suppressed in presence of bile. This suggests a hierarchical regulation of metabolism in bile stress. Among the electron acceptors generated by inflammatory host responses, nitrate has the highest standard redox potential (the E° for nitrate/nitrite redox couple is 433 mV), thus is energetically most favorable (19). Nitrate metabolism also suppresses anaerobic respiration dependent on other energetically inferior electron acceptors such as fumarate, tetrathionate and dimethyl sulfoxide (22, 29, 33). Tetrathionate acts as the respiratory electron acceptor that enables bacteria to utilize ethanolamine (32).

**Fig 8.**
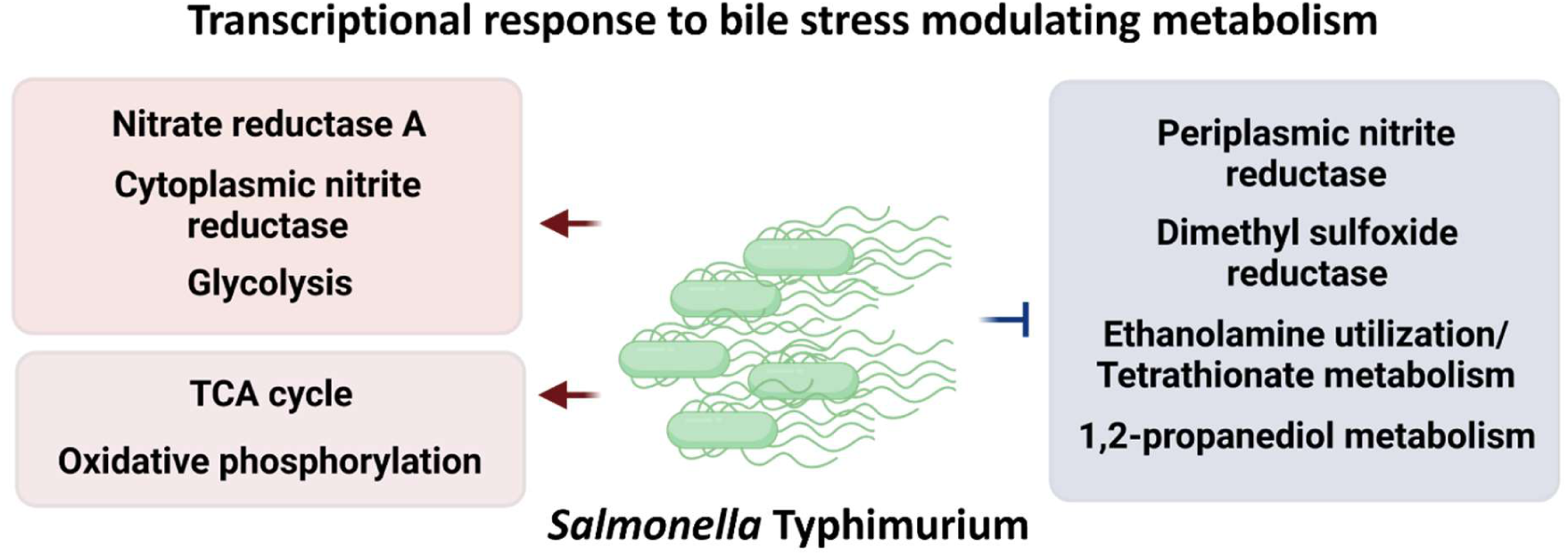
Illustration demonstrating *Salmonella* Typhimurium transcriptional response to bile stress. Modulation of metabolic processes involves induction of anaerobic respiration dependent on energetically superior electron acceptor nitrate while repressing energetically inferior processes that utilize dimethyl sulfoxide, tetrathionate and 1,2-propanediol.

While Fnr regulates the expression of target genes by sensing the oxygen levels, ArcA responds to the changes in levels of intracellular metabolites (40, 41). ArcA positively regulates the ethanolamine (*eut*) operon and the propanediol (*pdu*) operon (35). Although the *pdu* operon was suppressed in presence of bile, the downregulation was conspicuously higher in bile-sensitive Δ*csp*E than WT. A similar trend was observed in the transcript levels of dimethyl sulfoxide reductase and the ethanolamine utilization genes. Subsequently, we found that the Δ*arc*A strain was sensitive to bile stress (Fig 5), confirming the role of anaerobic respiration that is distinct to nitrate metabolism in mediating bile stress response. Taken together, these results indicate that even as nitrate-dependent anaerobic metabolism predominates adaptation to bile stress, other modes of anaerobic respiration may also be important in their basal levels and needs to be studied in future.

A possible explanation for the phenotype observed for Δ*fnr* and Δ*arc*A in bile stress is their protective role from reactive oxygen and nitrogen intermediates as bile is known to induce ROS (36, 37). The significant reduction of intracellular ROS observed in bile-sensitive Δ*csp*E strain upon overexpression of *fnr* and *arc*A demonstrates a mechanistic explanation for the above rationale (Fig 6C). Overall, our study reveals the substantial association of nitrate-dependent anaerobic metabolism and bile stress response in *S*. Typhimurium. The metabolic adaptation of *S*. Typhimurium to bile stress appears to mimic metabolism in inflamed gut. *S*. Typhimurium adopts anaerobic growth behavior in the presence of bile stress in aerated condition, possibly utilizing different carbon sources and electron acceptors to fulfill its energy requirements. The fact that bile-tolerant WT *S*. Typhimurium is better able to utilize anaerobic metabolism than bile-sensitive Δ*csp*E strain supports the significance of anaerobic metabolism. Also, our study presents the functional significance of *fnr* and *arc*A, important for nitrate and anaerobic metabolism, in mediating bile stress response.

## Material and Methods

### Bacterial strains and growth conditions

*Salmonella enterica* serovar Typhimurium ATCC 14028s was used as the WT strain and parent strain for all gene knockouts. Bacterial cultures were grown in Luria-Bertani (LB) medium at 37°C with aeration at 180 rpm. Single colony was inoculated, and the overnight grown culture was used as pre-inoculum for experiments. Ampicillin was added at the concentration of 100 μg/mL, when required. pQE60^ampR^ was used to overexpress *fnr* and *arc*A.

### Gene knockout preparation and cloning

Gene knockout strains were prepared by λ-red recombinase method described by Datsenko and Wanner (42). Briefly, pKD46 transformed *S*. Typhimurium strains were induced for recombinase expression using 20 mM L-arabinose (Sigma, USA). Cells were transformed with gene-specific linear DNA and selected on Kanamycin (50 µg/mL) (USP, VWR Life Science, USA) plates for gene deletion. Kanamycin cassette was removed by pcp20 transformation. For cloning, PCR amplification of *fnr* and *arc*A was performed using the WT genomic DNA as template. PCR amplified target genes and pQE60 plasmid (containing an IPTG inducible T5 promoter) were digested with *Bam*HI and *Hin*DIII (New England Biolabs) and purified by gel extraction (MinElute gel extraction kit, Qiagen, Germany). The purified insert was ligated into the cut pQE60 using T4 DNA ligase (Thermo Fisher Scientific) at 16°C overnight. *fnr* and *arc*A constructs were transformed into Top10 competent cells. p*fnr* and p*arc*A were purified (GeneJET plasmid miniprep kit, Thermo Fisher Scientific, Lithuania) and confirmed by visualization of insert release following restriction digestion and Sanger sequencing (Aggrigenome, Kerala, India). The strains and plasmids used in this study are listed in supplementary table S1. The knockout and cloning primers are listed in supplementary tables S2 and S3, respectively.

### RNA isolation

50 mL bacterial cultures were grown till O.D._600_ of 0.3 and then were treated with 3% bile for 90 minutes. Cultures were centrifuged and pellet was resuspended in 1 mL TRIzol reagent (Ambion, Invitrogen, USA) and allowed to lyse for 40 minutes at 1500 rpm in a shaking drybath (DLAB). 250 µL of chloroform (Sigma-Aldrich, USA) was added and mixed vigorously for 30 s. The samples were allowed to stand for 2 minutes and then centrifuged at 12,000 x g, 4 °C for 15 minutes. The aqueous phase was collected and mixed with 200 µL of chloroform and phase separation was repeated as earlier. Finally, 300 µL of aqueous phase was added to 500 µL of 2-propanol (Merck, Germany) and kept overnight at −20°C. The RNA was pelleted at 15,000 x g, 4 °C for 30 minutes and then washed with 75% ethanol, two times. Pellet was kept for air drying to remove residual ethanol and dissolved in nuclease-free water. RNA integrity was analysed on a 1.5% agarose gel by electrophoresis. RNA concentration and purity were determined using Nano-Drop (Thermo Fisher Scientific).

### cDNA library preparation and deep sequencing of the transcriptome

For RNA-Sequencing, the concentration of the extracted RNA samples was determined using a Nanodrop system (Nanodrop, Madison, USA), and the integrity of the RNA was examined by the RNA integrity number (RIN) using an Agilent 2100 bioanalyzer (Agilent, USA). In this experiment, each sample was set in triplicate. Further, rRNA depletion of the total RNA was done using RiboMinus kit (Thermo Fischer). Purified mRNA was fragmented into small pieces with fragment buffer at the appropriate temperature. Then, first-strand cDNA was generated using random hexamer-primed reverse transcription, followed by second-strand cDNA synthesis. Afterwards, A-Tailing Mix and RNA Index Adapters were added by incubating to end-repair the cDNA. The cDNA fragments obtained from the previous step were amplified by PCR, and the products were purified by Ampure XP Beads and then dissolved in EB solution. The product was validated on an Agilent Technologies 2100 bioanalyzer for quality control. Thus, prepared cDNA library was sequenced using Illumina NovoSeq 6000 platform with 150 bp read length in pair (150×2) yielding an average of 20 million reads per library.

### Quantitative and Qualitative resolution of expressed transcriptome

Raw data obtained in FASTq format was subjected to critical quality control of sequencing errors using FASTQC tool kit (http://www.bioinformatics.babraham.ac.uk/projects/fastqc) (43). Sequence reads which passed the Q30, after removing potential adapter Contamination were provided as input to Rockhopper (https://cs.wellesley.edu/~btjaden/Rockhopper/) (44, 45). *Salmonella* enterica subsp. enterica serovar Typhimurium str. LT2 was used as reference genome and paired sequence reads were aligned to identify potentially expressed transcripts spanning the whole genome. The output of rockhopper obtained as read count along with genomic co-ordinates and gene annotation was used as input to iGEAK software (46) for normalization and quality control of replicates. Upon quality control of the replicate reproducibility, differential gene expression analysis (DEGs) was performed with a minimum of 2-fold change and an FDR value of <0.05. Further, the differentially expressed genes were analysed for their functional roles and pathways enriched using PATRIC DB (https://www.bv-brc.org/app/Expression) (47). Various tools including Cluster 3.0 (http://eisenlab.org/), Java Treeview (48) and SRplot (https://www.bioinformatics.com.cn/srplot) was used to visualize the results and expression datasets.

### qRT-PCR

DNase I (New England Biolabs) treated RNA was used for cDNA preparation. 2.5 μg of RNA was reverse transcribed using RevertAid cDNA synthesis kit (Thermo Fisher Scientific, Lithuania). PCR reactions were prepared using 100 ng of cDNA samples, 250 nM of gene specific primers and SYBR green master mix (BioRad) and PCR was done in a Bio Rad CFX connect instrument. The primers (Sigma-Aldrich, Bangalore) for q-PCR are listed in supplementary tables S4 and S5. The normalized quantities of transcripts were estimated using the ΔΔCq method using two reference genes, *gmk* and *gyr*B. The WT grown in LB was used as control.

### Bacterial stress assays

Overnight grown cultures of indicated *S*. Typhimurium WT and gene knockout strains were diluted 1:500 in 50 mL LB followed by addition of bile (final concentration 3% v/v). For kinetics experiments, bile was added in the beginning. For complementation experiments, cells were grown to O.D._600_ of 0.3 followed by addition of bile. Cells were grown for 6 hours at 180 rpm, 37°C. 200 μL of culture from each growth condition was taken in clear flat bottom 96-well plate (Tarsons, Korea) and O.D._600_ was measured using Infinite 200-Pro instrument (Tecan, Austria GmbH). For CFU, appropriate dilutions were plated and grown at 37°C.

### ROS quantitation

Intracellular ROS quantitation was done as described previously (49). *S.* Typhimurium WT and Δ*csp*E strains containing vector (pQE60), p*fnr* or p*arc*A were diluted 1:250 in 5 mL LB and grown to O.D._600_ of 0.3 and treated with 3% bile for 6 hours. Cells were centrifuged and washed with 1X PBS to remove residual bile. Afterwards, cells were incubated with 20 µM 2’,7’-Dichlorofluorescin Diacetate (DCFDA) dye (Millipore, China) at 37°C for 30 minutes. Cells were washed twice with 1X PBS to remove the dye and finally resuspended in 300 μL of 1X PBS. 200 μL of cell suspension from each condition was taken in a 96-well plate. Fluorescence was recorded at excitation and emission wavelengths of 485 nm 535 nm respectively using Infinite 200-Pro instrument (Tecan, Austria GmbH). The values obtained were normalized to O.D._600_.

### Bile tolerance in presence of sodium nitrate

Overnight grown *S*. Typhimurium WT and Δ*csp*E strains were diluted 1:250 in LB with or without 30 mM Sodium nitrate. Cultures were incubated at 37°C, 180 rpm, grown to O.D._600_ of 0.2 and then treated with bile (Sigma-Aldrich) (final concentration 3% v/v) for 6 hours. O.D._600_ was measured, and cells were plated on LB agar at appropriate dilutions to count CFUs.

### Statistical analysis

Data analysis was done using GraphPad prism software. Two-way ANOVA was used to analyze the bacterial growth with each strain and condition compared using Tukey’s multiple comparisons test. One way ANOVA was used to analyze q-PCR data.

## Acknowledgements

We thank Varun Gowda and Madavan Vasudevan, Theomics International Pvt Ltd, Bengaluru for RNA sequencing and data analysis. We thank Siddharth Jhunjhunwala, BSSE, IISc for providing access to Biorender.

## Funding

This work was supported by core grant from IISc and the DBT-IISc partnership program. M.S. was supported by a fellowship from the Council of Scientific and Industrial Research, India.

## Competing interests

The authors declare no competing interests

## Notes

### Competing Interest Statement

The authors have declared no competing interest.

