## Supplemental Information for "Global transcriptome analysis reveals *Salmonella* Typhimurium employs the nitrate-dependent anaerobic pathway to combat bile stress"

**Supplementary tables:**

| **Table S1:** List of strains and plasmids Strain | Genotype/Features | Source/Reference |
|---|---|---|
| *S*. Typhimurium ATCC 14028s | Wild type, parent of all *Salmonella* strains | Ray *et al*., 2019 |
| *E.coli* TOP10 | F– *mcr*A Δ(*mrr*-*hsd*RMS-*mcr*BC) φ80*lac*ZΔM15 Δ*lac*X74 *rec*A1 *ara*D139 Δ(*ara-leu*)7697 *gal*U*gal*K λ– *rps*L(StrR) *end*A1 *nup*G | ThermoFisher Scientific |
| Δ*csp*E | 14028s; *csp*E::FRT | Ray *et al*., 2019 |
| Δ*nar*L | 14028s; *nar*L::FRT | This study |
| Δ*fnr* | 14028s; *fnr*::FRT | This study |
| Δ*arc*A | 14028s; *arc*A::FRT | This study |
| WT/VA | 14028s; pQE60; $Amp^r$ | This study |
| Δ*csp*E/VA | 14028s; pQE60; $Amp^r$ | This study |
| WT/p*fnr* | 14028s; pQE60; $Amp^r$ | This study |
| Δ*fnr*/ p*fnr* | 14028s; pQE60; $Amp^r$ | This study |
| Δ*csp*E/p*fnr* | 14028s; pQE60; $Amp^r$ | This study |
| WT/p*arc*A | 14028s; pQE60; $Amp^r$ | This study |
| Δ*arc*A/ p*arc*A | 14028s; pQE60; $Amp^r$ | This study |
| Δ*csp*E/p*arc*A | 14028s; pQE60; $Amp^r$ | This study |

**Table S2:** Primers used for gene knockout:

| Gene | Sequence 5'-3' |
|---|---|
| *nar*L FP | CACAGAAACCCAGGGAGATACCCATGAATAATCAG GAATAAGTGTAGGCTGGAGCTGCTTC |
| *nar*L RP | CGATAATCCGCATTGGCAACCGTTCCAGGAGCAAT AATTAGAATATCCTCCTTAGTTCCT |
| *fnr* FP | GTTAAAATTGACAAATATCAATTACGGCTTGAGCAG ACCTGTGTAGGCTGGAGCTGCTTC |
| *fnr* RP | ACGATATGGCAGAAGATAACATCAATGGTTTAGCTG ACGTTTAGAATATCCTCCTTAGTTCCT |
| *arc*A FP | ACTTCCTGTTTCGATTTAGTTGGCAATTTAGGTAGC AAACGTGTAGGCTGGAGCTGCTTC |
| *arc*A RP | TTAATCCTGCAGGTCGCCGCAGAAGCGATAACCTT CGGAATATCCTCCTTAGTTCCT |

**Table S3:** Primers used for cloning.

| Gene | Sequence 5'-3' |
|---|---|
| *fnr* FP | CGCGGATCCATGATCCCGGAAAAGCGAAT |
| *fnr* RP | CCCAAGCTTTTAAGCGACGTTGCGGGTATG |
| *arc*A FP | CGCGGATCCATGCAGACCCCGCACATTCT |
| *arc*A RP | CCCAAGCTTTTAATCCTGCAGGTCGCCGC |

**Table S4:** Primers used for validation of RNA-seq data using q-PCR.

| Gene | Sequence 5’-3’ |
|---|---|
| *ygb*L FP | TGACGATACTGGCAAAAGAA |
| *ygb*L RP | GAGATTACCCAGACACGAAC |
| *nar*K FP | CTGTCTAAAACGCAATTCCC |
| *nar*K RP | AGGACAAAGTTGATAAGCGT |
| *cyo*A FP | ACTCAGTGGCTGTAATTCTG |
| *cyo*A RP | ATTACTCGCACGATACTTCC |
| *cyo*B FP | AACCTATCGTCATGGTAACG |
| *cyo*B RP | CACGTAACAGCATCACAATC |
| *cyo*D FP | TGACGGTTATTCCATTCTGG |
| *cyo*D RP | ATCAGCACGGTAAAGATGAA |
| *pdu*A FP | GCTATGAAAAGATTGGCTCC |
| *pdu*A RP | GCTAATTCCCTTCGGTAAGA |
| *omp*W FP | GATTACCTGATTAACCGCGA |
| *omp*W RP | GCCGAGAACATAAATACCCA |
| *yde*Y FP | TTTATTGCTGGCCTGGTATT |
| *yde*Y RP | GTTTTTGCTTCCGTCTGTAG |
| *yna*F FP | CTGAAGCAAAGATTGACGAC |
| *yna*F RP | ATCGCTTCCAGTTGAGATTT |
| *yne*B FP | GGCAATCAATAAACCAGTCG |
| *yne*B RP | GATGTTCATGCTCACTACCA |

**Table S5:** Primers used for q-PCR:

| Gene: | Sequence 5’-3’ |
|---|---|
| *nar*L FP | AGGACGTATTGTCGTTTTCA |
| *nar*L RP | CACTTAATACCATTTCGCCG |
| *nar*G FP | GAAAGTTTGCTGGTGTATCG |
| *nar*G RP | CAGCATCAACAGATTATCGC |
| *nar*H FP | GGTGCTGAATCTCGATAAGT |
| *nar*H RP | CAGTCATTCGGAAAACCAAC |
| *nar*I FP | TGGTTGGCGGTATTTTACTG |
| *nar*I RP | TCATTTCACTCCCGTCCATA |
| *nir*D FP | TACCATAGCGATCAGGTGTT |
| *nir*D RP | ATGCATAAACCATCGCTCAG |
| *nrf*B FP | AAGGTATGCACGGTAAACAT |
| *nrf*B RP | CTGCTCTACCGTATACATCG |
| *fnr* FP | TTAACGAGCATGAGCTTGAT |
| *fnr* RP | TGTAGCTCTTAATCGTTCCG |
| *arc*A FP | AAAAACGGTCTCCTGTTAGC |
| *arc*A RP | TAGTTTATACTGCTCGCCG |
| *nar*P FP | AATATGAAAGGTCTGAGCGG |
| *nar*P RP | GATCGCTATCTTTGAGCAGA |
| *nap*H FP | CCGTATCAGCCAGTTTATGG |
| *nap*H RP | GACTTTCCAGGGTAATCAGC |

**Table S6: WT vs W**

| No. | Gene name/ ID | Description | $Log_2$ Fold Change | adj. p Val |
|---|---|---|---|---|
| 1. | *nar*K | nitrite extrusion protein | 7.28 | 0.00 |
| 2. | *ygb*L | putative aldolase | 6.26 | 0.00 |
| 3. | *ygb*K | putative tRNA synthase | 5.59 | 0.00 |
| 4. | STM14_0677 | hypothetical protein | 5.16 | 0.00 |
| 5. | *ygb*M | hypothetical protein | 5.09 | 0.00 |
| 6. | *nar*G | nitrate reductase 1 subunit alpha | 5.02 | 0.00 |
| 7. | *cyo*A | cytochrome o ubiquinol oxidase subunit II | 4.81 | 0.00 |
| 8. | *fad*B | multifunctional fatty acid oxidation complex subunit alpha | 4.78 | 0.00 |
| 9. | *nap*F | ferredoxin-type protein | 4.63 | 0.00 |
| 10. | *cyo*D | cytochrome o ubiquinol oxidase subunit IV | 4.55 | 0.00 |
| 11. | STM14_3516 | putative nucleoside-diphosphate-sugar epimerase | 4.48 | 0.00 |
| 12. | *cyo*B | cytochrome o ubiquinol oxidase subunit I | 4.34 | 0.00 |
| 13. | *rpl*V | 50S ribosomal protein L22 | 4.29 | 0.00 |
| 14. | *lld*P | L-lactate permease | 4.28 | 0.00 |
| 15. | *rps*S | 30S ribosomal protein S19 | 4.23 | 0.00 |
| 16. | *cyo*C | cytochrome o ubiquinol oxidase subunit III | 4.21 | 0.00 |
| 17. | *nap*D | assembly protein for periplasmic nitrate reductase | 4.17 | 0.00 |
| 18. | *cyo*E | protoheme IX farnesyltransferase | 4.14 | 0.00 |
| 19. | *lld*R | DNA-binding transcriptional repressor LldR | 4.08 | 0.00 |
| 20. | STM14_3932 | hypothetical protein | 4.03 | 0.00 |

| No. | Gene name/ ID | Description | $Log_2$ Fold Change | adj. p Val |
|---|---|---|---|---|
| 1. | *ygh*W | putative cytoplasmic protein | -5.35 | 0.00 |
| 2. | *pdu*A | polyhedral body protein | -5.01 | 0.00 |
| 3. | *pud*B | polyhedral body protein | -4.50 | 0.00 |
| 4. | *ssa*G | type III secretion system apparatus protein | -4.37 | 0.00 |
| 5. | *yne*B *lsr*F | aldolase | -4.09 | 0.00 |
| 6. | STM14_0894 | putative ABC transport protein | -4.08 | 0.00 |
| 7. | *pdu*K | polyhedral body protein | -4.01 | 0.00 |
| 8. | *omp*W | outer membrane protein W | -3.96 | 0.00 |
| 9. | *yna*F | putative universal stress protein | -3.91 | 0.00 |
| 10. | *yde*Y | putative sugar transport protein | -3.90 | 0.00 |
| 11. | *yde*Z | putative sugar transport protein | -3.81 | 0.00 |
| 12. | STM14_5185 | hypothetical protein | -3.72 | 0.00 |
| 13. | STM14_3806 | putative periplasmic ferrichrome-binding protein | -3.70 | 0.00 |
| 14. | *zra*P | zinc resistance protein | -3.64 | 0.00 |

| | | | | |
|---|---|---|---|---|
| 15. | *pdu*J | polyhedral body protein | -3.64 | 0.00 |
| 16. | *ssa*H | type III secretion system apparatus protein | -3.63 | 0.00 |
| 17. | STM14_1496 | hypothetical protein | -3.63 | 0.00 |
| 18. | *ssa*I | type III secretion system apparatus protein | -3.56 | 0.00 |
| 19. | *eut*M | putative detox protein | -3.46 | 0.00 |
| 20. | STM14_4895 | putative mannose-6-phosphate isomerase | -3.41 | 0.00 |

**Table S7: ET vs W**

| No. | Gene name/ ID | Description | $Log_2$ Fold Change | adj. p Val |
|---|---|---|---|---|
| 1. | *ygb*L | putative aldolase | 6.81 | 0.00 |
| 2. | *nar*K | nitrite extrusion protein | 6.63 | 0.00 |
| 3. | *ygb*K | putative tRNA synthase | 6.11 | 0.00 |
| 4. | *ygb*M | hypothetical protein | 5.65 | 0.00 |
| 5. | *cyo*A | cytochrome o ubiquinol oxidase subunit II | 5.54 | 0.00 |
| 6. | *glp*D | glycerol-3-phosphate dehydrogenase | 5.08 | 0.00 |
| 7. | *cyo*B | cytochrome o ubiquinol oxidase subunit I | 4.96 | 0.00 |
| 8. | *rpl*V | 50S ribosomal protein L22 | 4.95 | 0.00 |
| 9. | *rps*S | 30S ribosomal protein S19 | 4.90 | 0.00 |
| 10. | STM14_0296 | hypothetical protein | 4.89 | 0.00 |
| 11. | STM14_4797 | hypothetical protein | 4.89 | 0.00 |
| 12. | STM14_0677 | hypothetical protein | 4.81 | 0.00 |
| 13. | *glp*A | sn-glycerol-3-phosphate dehydrogenase subunit A | 4.78 | 0.00 |
| 14. | STM14_4970 | hypothetical protein | 4.73 | 0.00 |
| 15. | *ygb*J | 3-hydroxyisobutyrate dehydrogenase | 4.71 | 0.00 |
| 16. | *rpl*W | 50S ribosomal protein L23 | 4.61 | 0.00 |
| 17. | *pri*B | primosomal replication protein N | 4.60 | 0.00 |
| 18. | *glp*T | sn-glycerol-3-phosphate transporter | 4.56 | 0.00 |
| 19. | STM14_3516 | putative nucleoside-diphosphate-sugar epimerase | 4.56 | 0.00 |
| 20. | *cyo*D | cytochrome o ubiquinol oxidase subunit IV | 4.55 | 0.00 |

| No. | Gene name/ ID | Description | $Log_2$ Fold Change | adj. p Val |
|---|---|---|---|---|
| 1. | *pdu*A | polyhedral body protein | -6.52 | 0.00 |
| 2. | *flj*B | flagellin | -5.99 | 0.00 |
| 3. | *ygh*W | putative cytoplasmic protein | -5.61 | 0.00 |
| 4. | predicted RNA_3753 | predicted RNA | -5.22 | 0.00 |
| 5. | *yde*Y | putative sugar transport protein | -5.21 | 0.00 |
| 6. | STM14_3799 | putative methyl-accepting chemotaxis protein | -5.20 | 0.00 |
| 7. | *yde*Z | putative sugar transport protein | -4.98 | 0.00 |
| 8. | *yne*B | aldolase | -4.94 | 0.00 |
| 9. | *lsr*A | putative ABC-type aldose transport system ATPase component | -4.80 | 0.00 |
| 10. | *yna*F | putative universal stress protein | -4.73 | 0.00 |
| 11. | *omp*W | outer membrane protein W | -4.71 | 0.00 |
| 12. | STM14_2852 | putative chemotaxis signal transduction protein | -4.69 | 0.00 |
| 13. | *pud*B | polyhedral body protein | -4.56 | 0.00 |
| 14. | *flg*M | anti-sigma-28 factor FlgM | -4.54 | 0.00 |

| | | | | |
|---|---|---|---|---|
| 15. | *yci*G | putative cytoplasmic protein | -4.46 | 0.00 |
| 16. | STM14_4895 | putative mannose-6-phosphate isomerase | -4.40 | 0.00 |
| 17. | STM14_1578 | hypothetical protein | -4.30 | 0.00 |
| 18. | *yne*A | putative sugar transport protein | -4.25 | 0.00 |
| 19. | *tcp* | methyl-accepting transmembrane citrate/phenol chemoreceptor | -4.25 | 0.00 |
| 20. | STM14_1579 | pyrimidine (deoxy)nucleoside triphosphate pyrophosphohydrolase | -4.23 | 0.00 |

**Supplementary figures:**

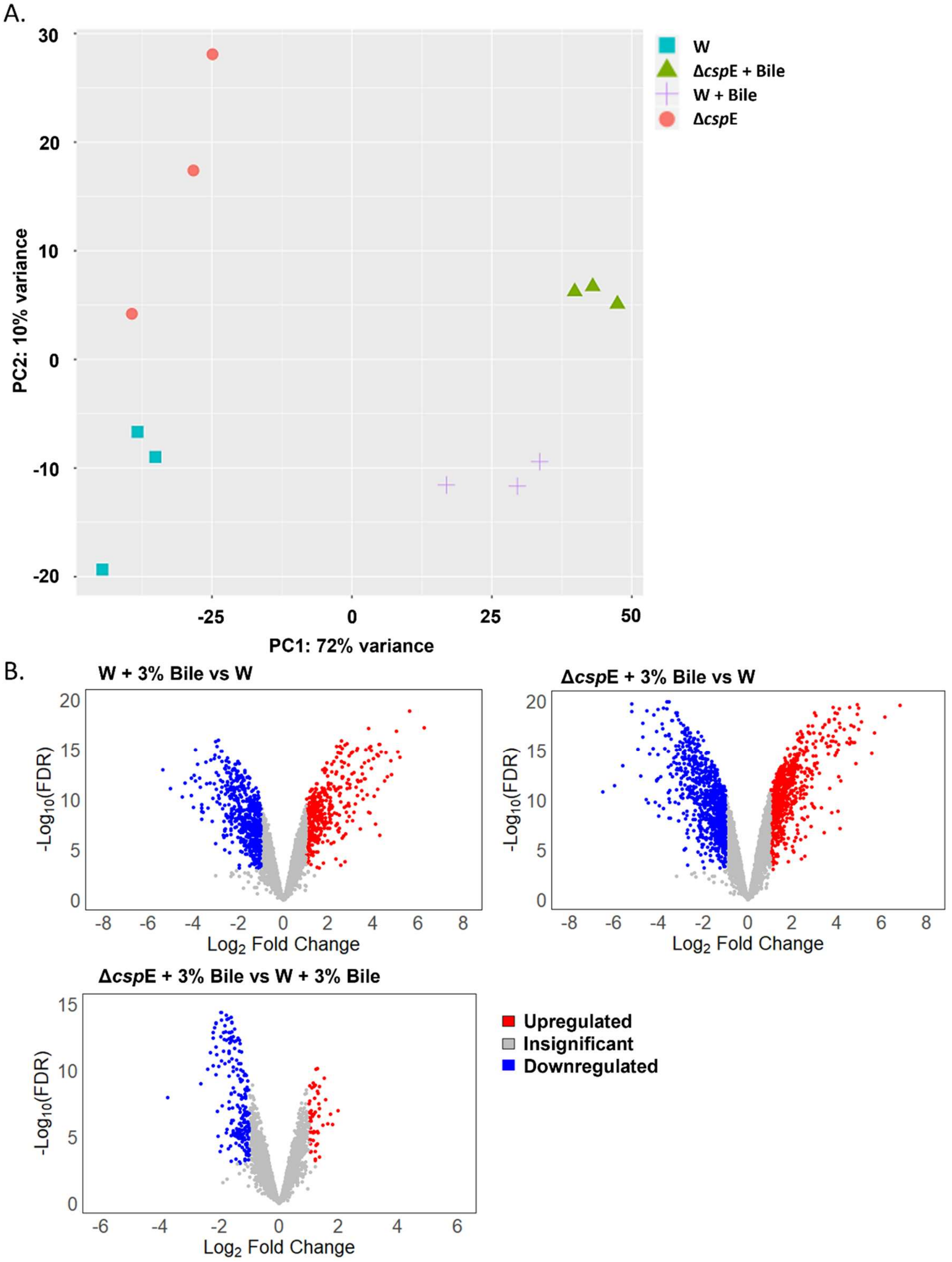


**Fig S1:** A. Principal component analysis of control and bile treated WT and Δ*csp*E strains. B. Volcano plots depicting statistically significant changes in transcripts levels upon bile treatment.

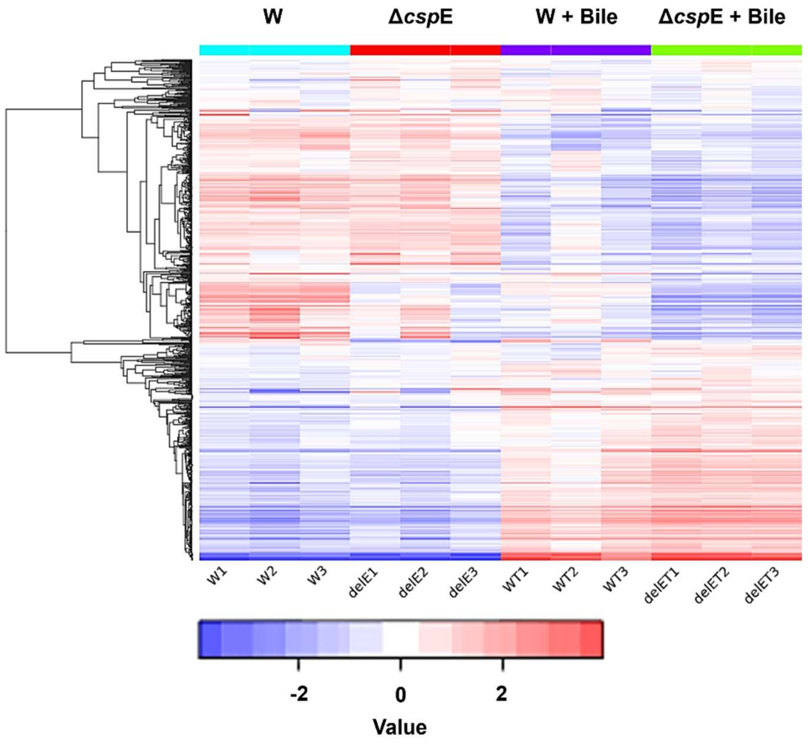


**Fig S2:** Heatmap demonstrates a distinct pattern of differentially expressed upregulated and downregulated genes in control and bile treated WT and Δ*csp*E strains.

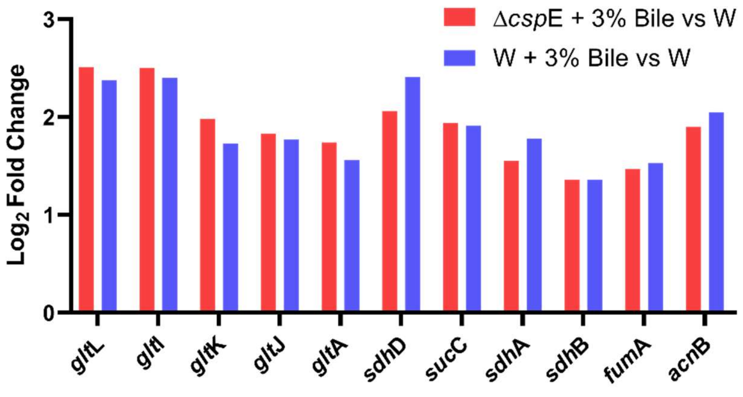


**Fig S3:** Histogram showing the differential expression of genes involved in Citrate/TCA cycle in bile treated WT and Δ*csp*E strain as $Log_2$ fold change values.

A.

| Genes | WT (3% Bile) vs WT (LB) ($Log_2$ Fold Change) | Δ*csp*E (3% Bile) vs W (LB) ($Log_2$ Fold Change) |
|---|---|---|
| *nar*K | 7.28 | 6.63 |
| *ygb*L | 6.26 | 6.81 |
| *cyo*A | 4.81 | 5.54 |
| *cyo*B | 4.34 | 4.96 |
| *cyo*D | 4.55 | 4.55 |

B.

| Genes | WT (3% Bile) vs WT (LB) ($Log_2$ Fold Change) | Δ*csp*E (3% Bile) vs W (LB) ($Log_2$ Fold Change) |
|---|---|---|
| *pdu*A | -5.01 | -6.52 |
| *yne*B | -4.09 | -4.94 |
| *omp*W | -3.96 | -4.71 |
| *yna*F | -3.91 | -4.73 |
| *yde*Y | -3.90 | -5.21 |

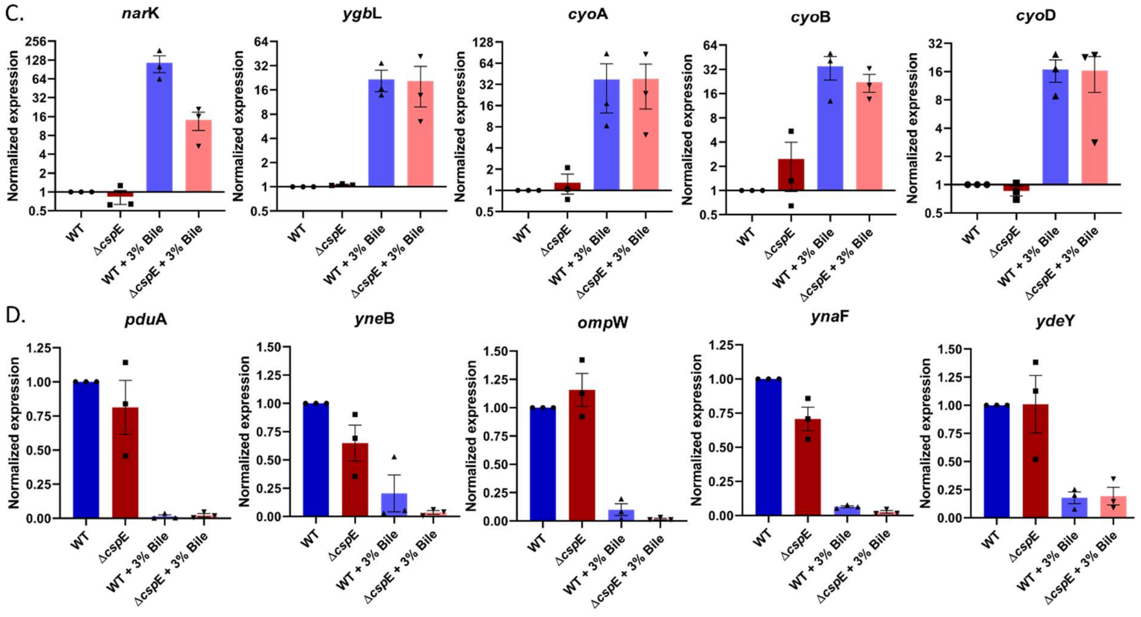


**Fig S4:** qRT-PCR validation of transcriptional changes detected in RNA-Seq, in response to bile involving alteration in metabolism and quorum sensing. *S.* Typhimurium WT and Δ*csp*E strains were grown to O.D. 0.3 and treated with 3% bile for 90 minutes. Expression of A. five most upregulated genes and B. five most downregulated genes common to bile-treated WT and Δ*csp*E strains in RNA-seq. C. Normalized expression of upregulated genes based on qRT-PCR data. D. qRT-PCR analysis of downregulated genes identified in RNA-seq. Data is representative of 3 independent experiments.

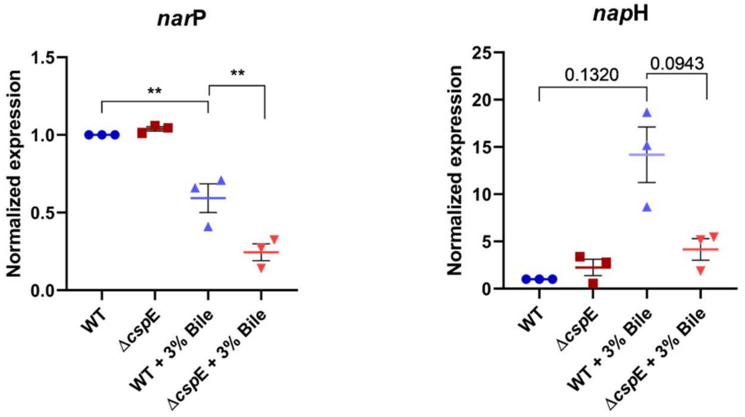


**Fig S5:** q-RT PCR analysis of genes involved in nitrate metabolism under low nitrate conditions following 3% bile treatment. *nar*P is response regulator of two-component system while *nap*H is required for electron transfer to the periplasmic nitrate reductase. Data shown as mean±SEM and is representative of 3 independent experiments. P values were measured by one-way ANOVA using Sidak's multiple comparisons test. **p<0.01.

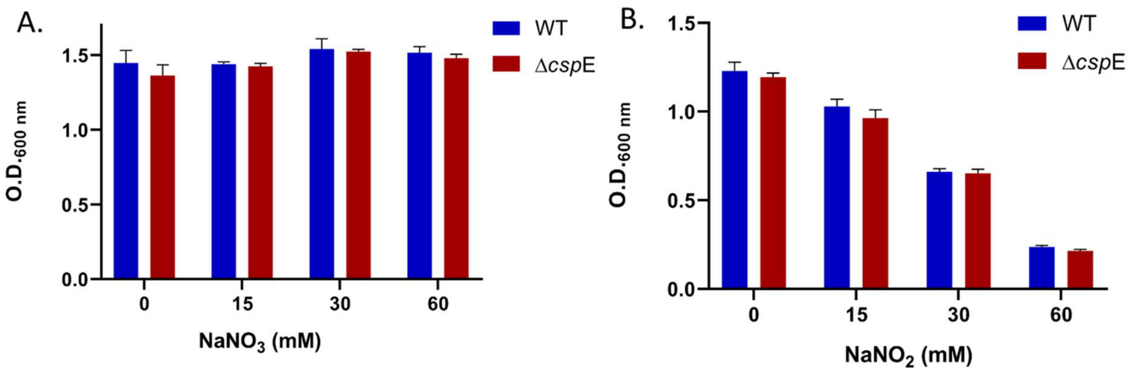


**Fig S6:** A. Growth of WT and Δ*csp*E strains in different concentrations of sodium nitrate. B. Growth of WT and Δ*csp*E in presence of different concentrations of sodium nitrite. Data shown as mean±SD and is representative of 3 independent experiments.

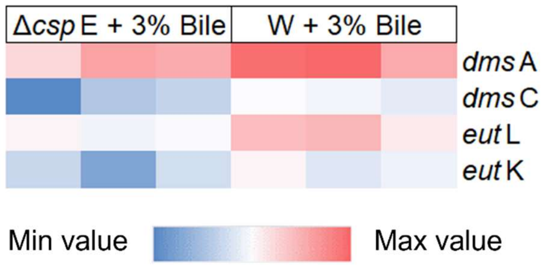


**Fig S7:** Heat map showing genes involved in anaerobic respiration and differentially expressed between bile treated Δ*csp*E and WT.
